# Transcription Factors STAT5A and SPI1 Reveals RHBDD2 as a Potential Biomarker in Sepsis and Septic Shock

**DOI:** 10.1101/2020.09.15.285551

**Authors:** Arslan Ali, Huma Shehwana, Ayesha Hanif, Abeera Fatima, Maria Shabbir, Mehak Rafiq

## Abstract

Sepsis is a serious health situation caused by uncontrolled infection and septic shock is a severe condition of sepsis. RHBDD2 is a member of the rhomboid superfamily which is overexpressed in different types of cancer and associated with ER stress and estrogen receptor. Using microarray gene expression data and using different computational techniques this study investigated the role of RHBDD2 in sepsis and septic shock. Finds functional annotation of RHBDD2 using co-expression analysis and identified the deregulation of RHBDD2 in sepsis using differential expression analysis. Results show that RHBDD2 is overexpressed in sepsis and septic shock. The GO enrichment analysis, KEGG pathways, and biological functions of the RHBDD2 co-expressed genes module show that it is involved in most of the sepsis-related biological functions and also plays a role in most of the infection-related pathways which lead to sepsis and septic shock. RHBDD2 is regulated by STAT5A and SPI1 transcription factors in sepsis and septic shock. The identification of the RHBDD2 as a biomarker may facilitate in septic shock diagnosis, treatment, and prognosis.

## Introduction

Sepsis is a condition that puts the body of an individual in the closed risk of death and its incidence is increasing day by day, although the death rate has lessened to a minor extent^1,2^. The treatment of sepsis is amended by the ages by using the early goal-directed method but the identification procedure and its biomarkers need to be worked on a lot thus far^3^. Persisting sepsis can result in progression which as a consequence leads to septic shock regarded as cellular, cardiovascular metabolic aberration^4^. Septic shock is even more life-threatening than simply sepsis; it is an exacerbated form of sepsis. Sepsis along with septic shock is a source of 37% to 56% under treatment deaths in the hospital ICUs. Every year, sepsis is estimated to affect no less than 30 million people worldwide resulting in 6 million deaths. Just like cancer sepsis is the most prevalent in low- and middle-income countries. It is estimated that worldwide 3 million newborn babies and 1.2 million children suffer from sepsis every year^5^. Sepsis was termed a global health priority by the World Health Organization (WHO) and the World Health Assembly (WHA), they also implemented a resolution to enforce the United Nation (UN) members to develop the diagnosis, prevention, and supervision of sepsis. This study like to find the role of RHBDD2 in sepsis and septic shock. RHBDD2 is a distantly related member of the Rhomboid superfamily of membrane-bound proteases that catalyze regulated intramembrane proteolysis which involved in different types of cancer. Most of the forgoing studies determine that RHBDD2 is associated with breast cancer and in advance stages of colorectal cancer^6^. Analysis of EGFR signaling in drosophila led to the discovery of Rhomboid-1 which is the 1st Rhomboid intermembrane serine protease a new class of enzyme^7^. The rhomboid proteases to exist in almost all prokaryotes and eukaryotes The prior studies show that RHBDD2 expression in Breast Cancer. RHBDD2 is a novel oncology-related biomarker that shows overexpression at mRNA and protein level in Breast cancer and plays as a facilitator in breast cancer progression. RHBDD2 expression has been related to proliferation, migration, and cellular response to ER stress^8^. Over-expression RHBDD2 has also a significant association among breast carcinomas with low-negative progesterone receptor expression^9^. RHBDD2 also involved in ER stress but its specific role is not cleared^10^. The earlier studies did not investigate the role of RHBDD2 in sepsis and septic shock. But in this research for the first time, we evaluate the association of RHBDD2 with sepsis and septic shock. The links to associate the Septic Shock with RHBDD2 are following

- ER stress
- Estrogen Receptor

RHBDD2 has involved by ER stress mention in Cancer biology^10^. ER stress also involved in Septic Shock and its former stage Sepsis plays a dynamic role and take part in the pathogenesis of Septic shock and Sepsis^11^. Research on septic rats discovered that constituents of ER stress (CHOP, caspase-12, and GRP94) were up-regulated in the hearts of septic rats, and the stoppage of the ER stress-induced apoptosis and defended the myocardial cells from ER stress^12^. Estrogen Receptor has an significant role in cancer and protective role in sepsis. Studies show that estrogen receptors constructively influence chemotaxis of neutrophils, cytokine release, Heat Shock Proteins(HSP) expression, Heme oxygenase 1 (HO-1) induction, and the refurbishment of organ functions in Septic shock and sepsis^13^. In the pathway selective Estrogen receptor ligands show a potential role in Septic Shock and Sepsis^14^. Inflammatory response reduced by Estrogen receptor *α* in the small intestine and liver while the Estrogen Receptor *β* up-regulated inflammatory response in the small intestine and lung^15^. ER Stress is implicated in sepsis alongside with estrogen receptor. In the cancer estrogen receptor is regulated by RHBDD2. However, it has not been studied yet in sepsis and septic shock even though it is a regulator for ER Stress in cancer. This study looks at the role of RHBDD2 in sepsis and septic shock.

## Results

### 0.1 Datasets selection and pre-processing

Seven Affymetrix datasets were the result of datasets selection criteria using different filtering threshold (see Fig). GSE4607, GSE8621, GSE9692, GSE11755, GSE26440, GSE26378 and GSE13904. Information about the datasets is shown in Table 1

**Table 1.**
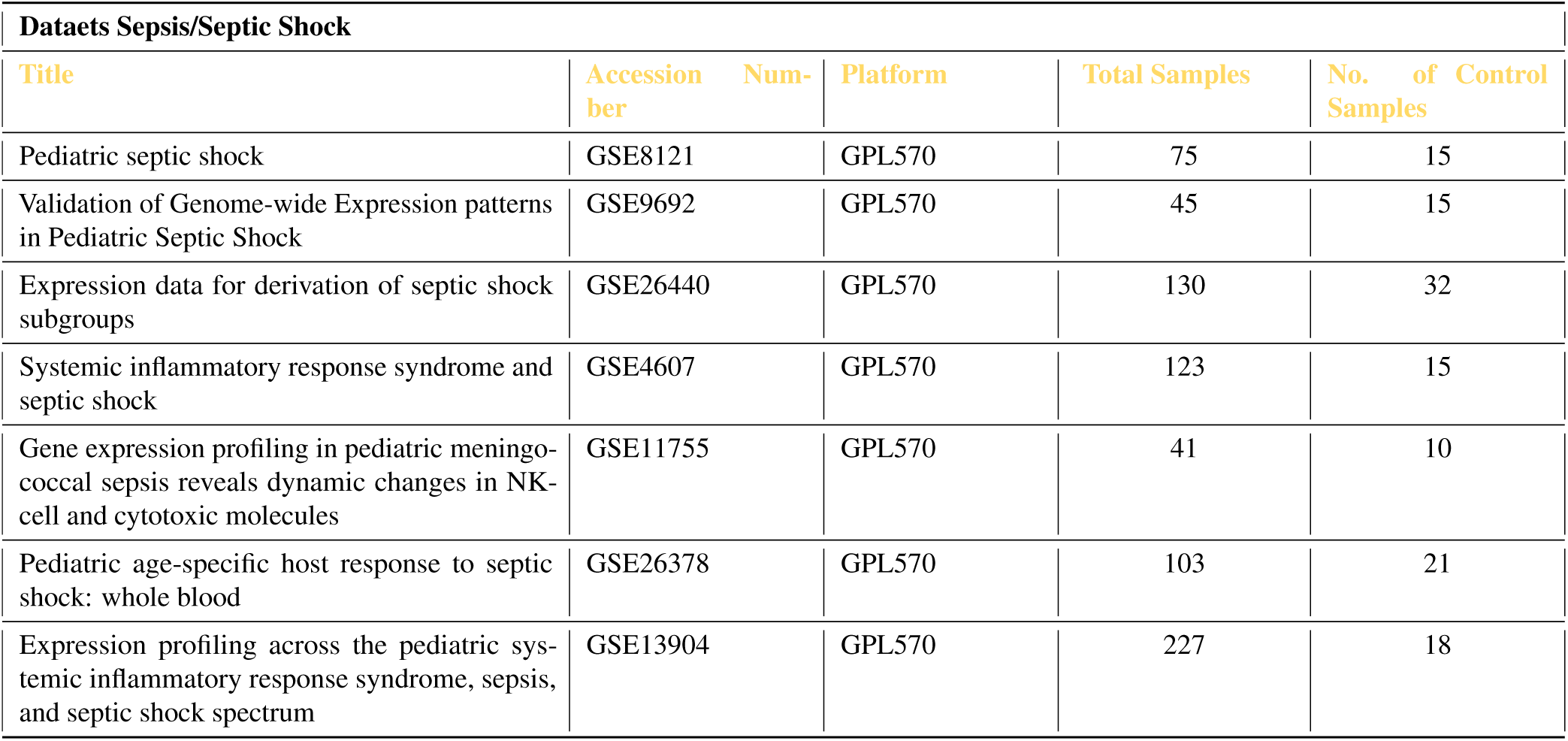
Datasets included in meta-analysis

After the quality check, no sample was identified as outlier in any of seven datasets. MAS5 detect 54675 genes in each dataset which remained 20517 for further analysis in each dataset after Jetset reduction criteria. Principle component analysis (PCA) plots of all seven datasets shows fine cluster to differentiate control and disease samples shown in Figure 6.

**Figure 1.**
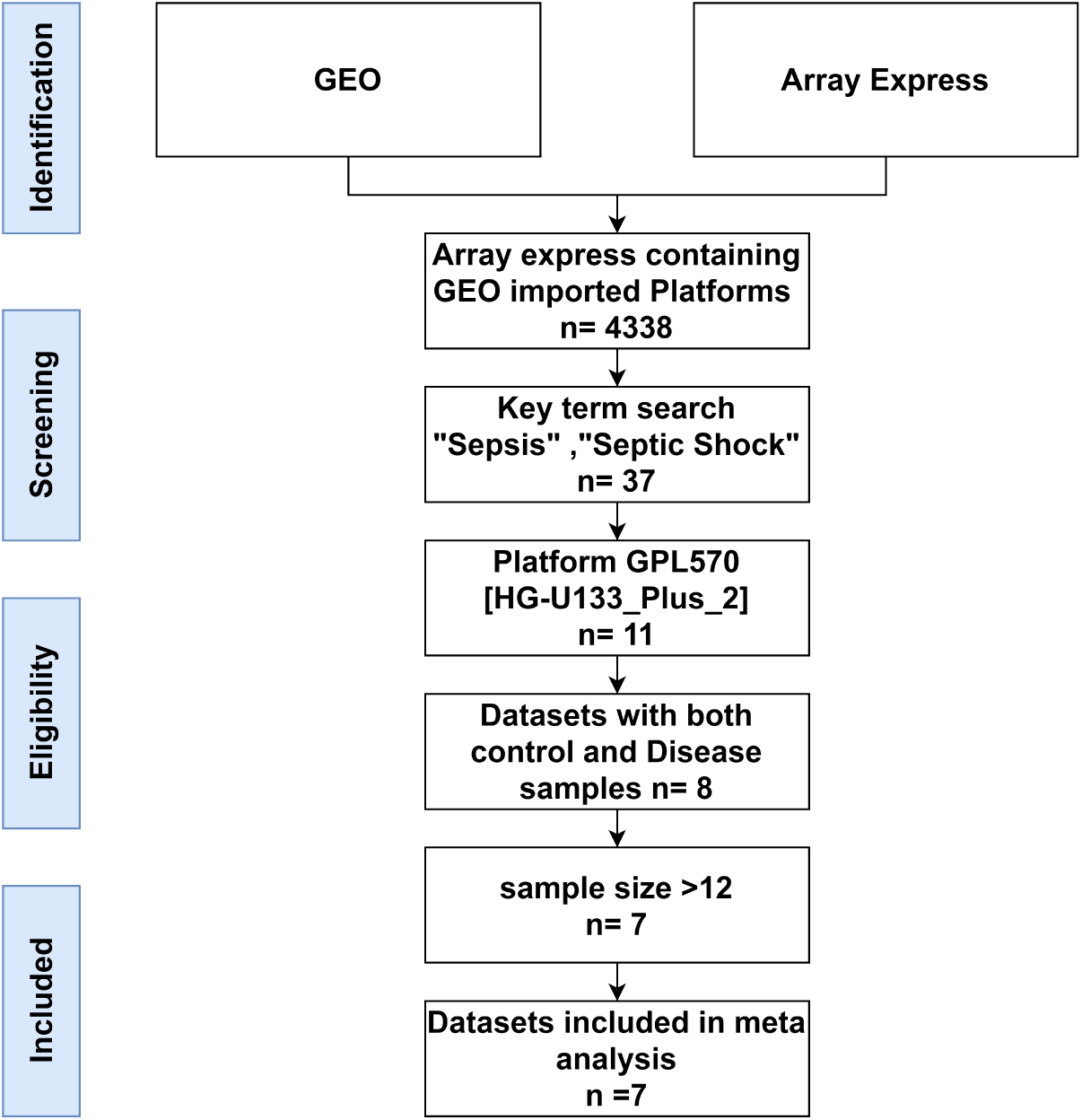
Flow diagram of datasets selection.

**Figure 2.**
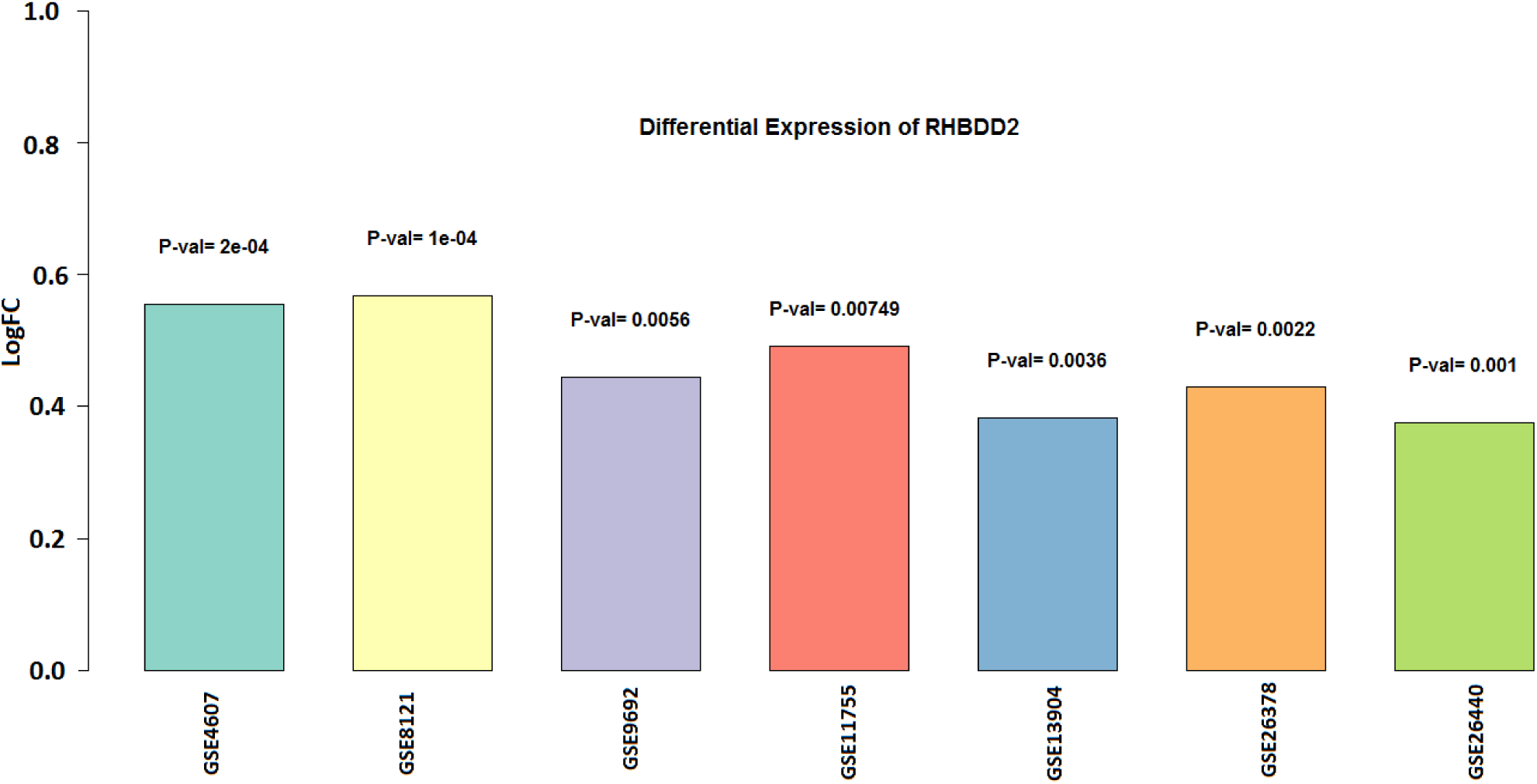
Bar plot of LogFC value of seven datasets with P-value showing on the head of each bar

**Figure 3.**
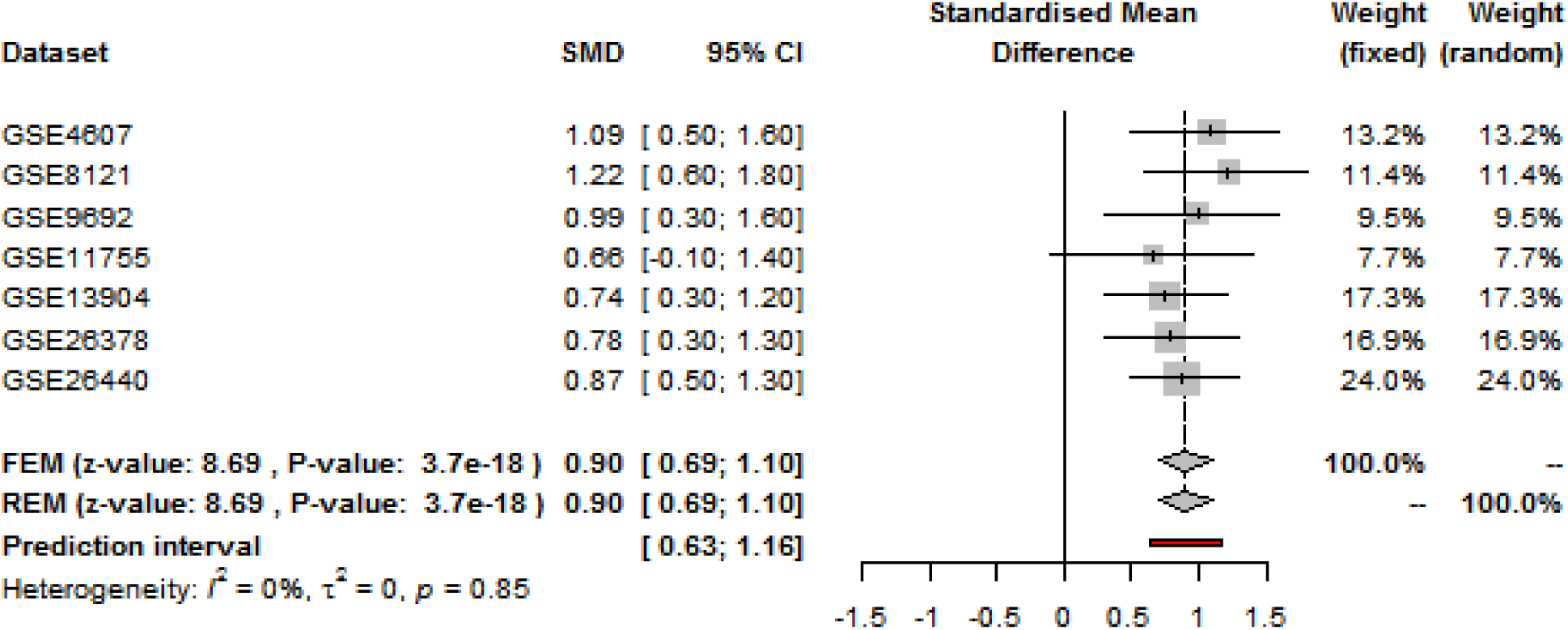
Forest plot of seven datasets of sepsis and septic shock showing the result of meta-analysis of differential expression of RHBBD2

**Figure 4.**
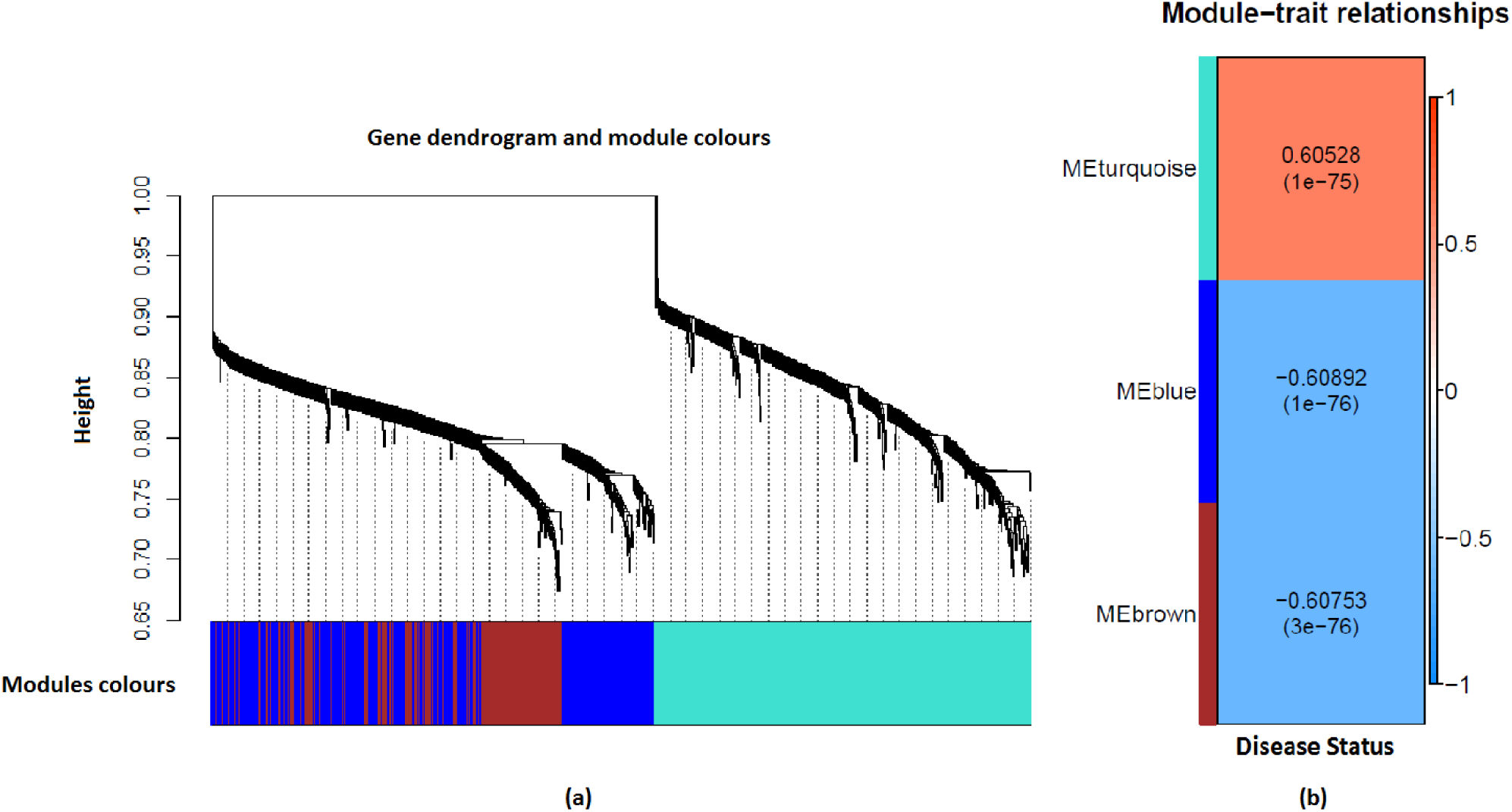
The clustering of gene in the form of modules and their relationship with the disease. (a) represents the clustering of 1000 correlated genes with RHBDD2 in form of modules using WGCNA. (b) represents the correlation of different modules with sepsis and septic shock and their significance in the form of P-value. The intensity of orange color shows positive correlation wherease the intensity of blue color shows the negative correlation.

**Figure 5.**
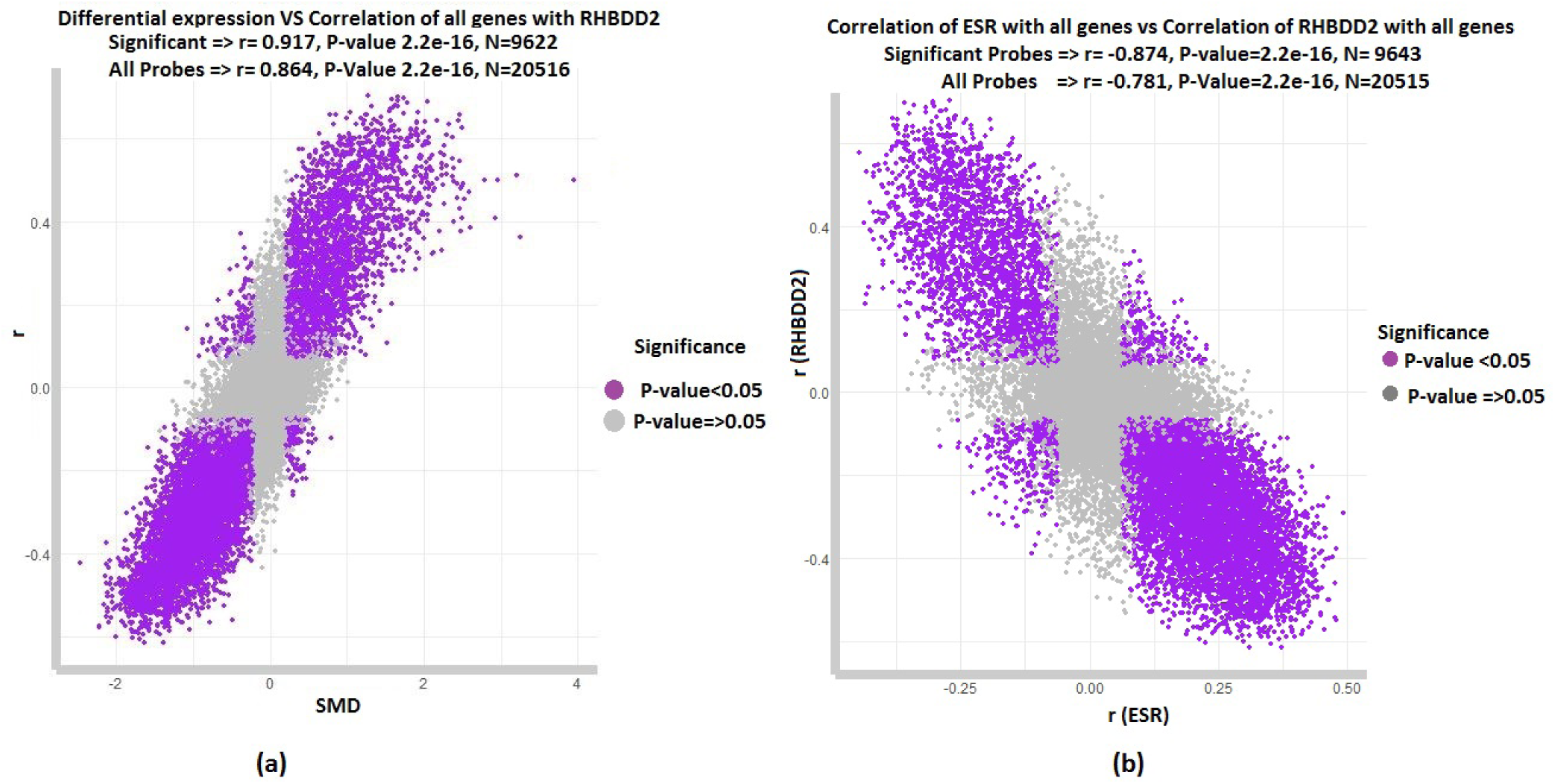
(a) graph showing the relationship of meta-differential expression and meta-correlation of all genes with RHBDD2 X-axis represent the standard mean difference obtained using GeneMeta and Y-axis representing the meta-correlation of all genes with RHBDD2. (b) Graph showing the relationship of ESR and RHBDD2 in sepsis septic shock X-axis represent the meta-correlation of ESR with all genes and Y-axis representing the meta-correlation of RHBDD2 with all genes in seven datasets of sepsis septic shock

**Figure 6.**
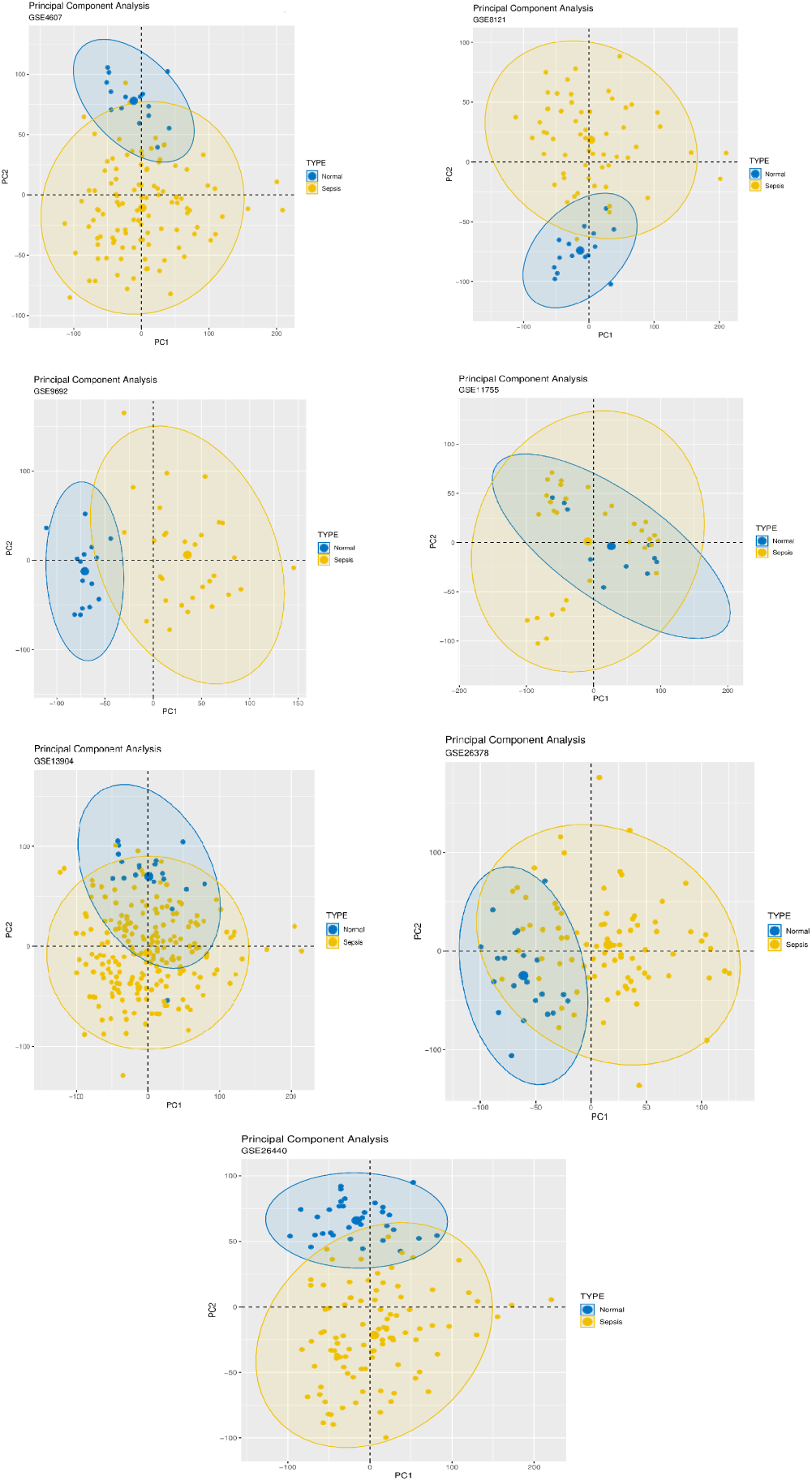
PCA plots of all seven datasets

### 0.2 Meta-Analysis

#### 0.2.1 Differential Expression Analysis

Microarray analysis revealed that RHBDD2 up-regulated gene in Sepsis and septic shock patients compared with the normal group patients. Limma Results in Figure2 presenting, RHBDD2 has significant logFC value in each dataset ranging (0.4-0.6) with Pvalue< 0.05. GeneMeta, the meta differential expression showing with FDR <0.05 and Muvals (>0.5 or less <-0.5) obtained 7377 genes as differential expression genes from which 2022 were over expressed and RHBDD2 ranked at 995 in over expressed genes and 5355 were down expressed.

The forest plot in Figure 3 showing that from both Random effect model(REM) and Fixed effect model (FEM) showed same results with pooled SMD (0.90)[0.69, 1.10] and with same FEM and REM (z-value of 8.69, P-value 3.7e-18) by using the both fixed and random weights which mean that RHBDD2 expression is significantly is higher in sepsis and septic shock as compare to normal individual. Over all 95% CI lies between [0.63, 1.16]. Heterogeneity is 0% which with P-value(0.85) showing consistency between study.

#### 0.2.2 Co-expression Analysis

WGCNA analysis acquired three modules from 1000 genes which were top correlated genes with RHBDD2 shown in Figure 4. Their relation with disease status was found which showed that module turquoise carried the genes which were positively correlated with disease and RHBDD2. While blue and brown carried genes which were negatively correlated with RHBDD2 and were down regulated in disease. The distribution of WGCNA modules is shown in Table 2

**Table 2.**
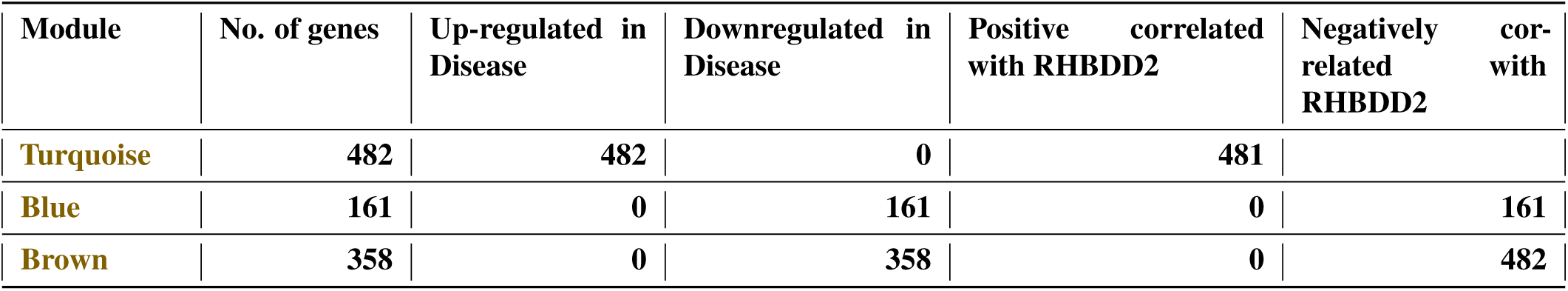
Distribution of WGCNA modules

**Table 3.**
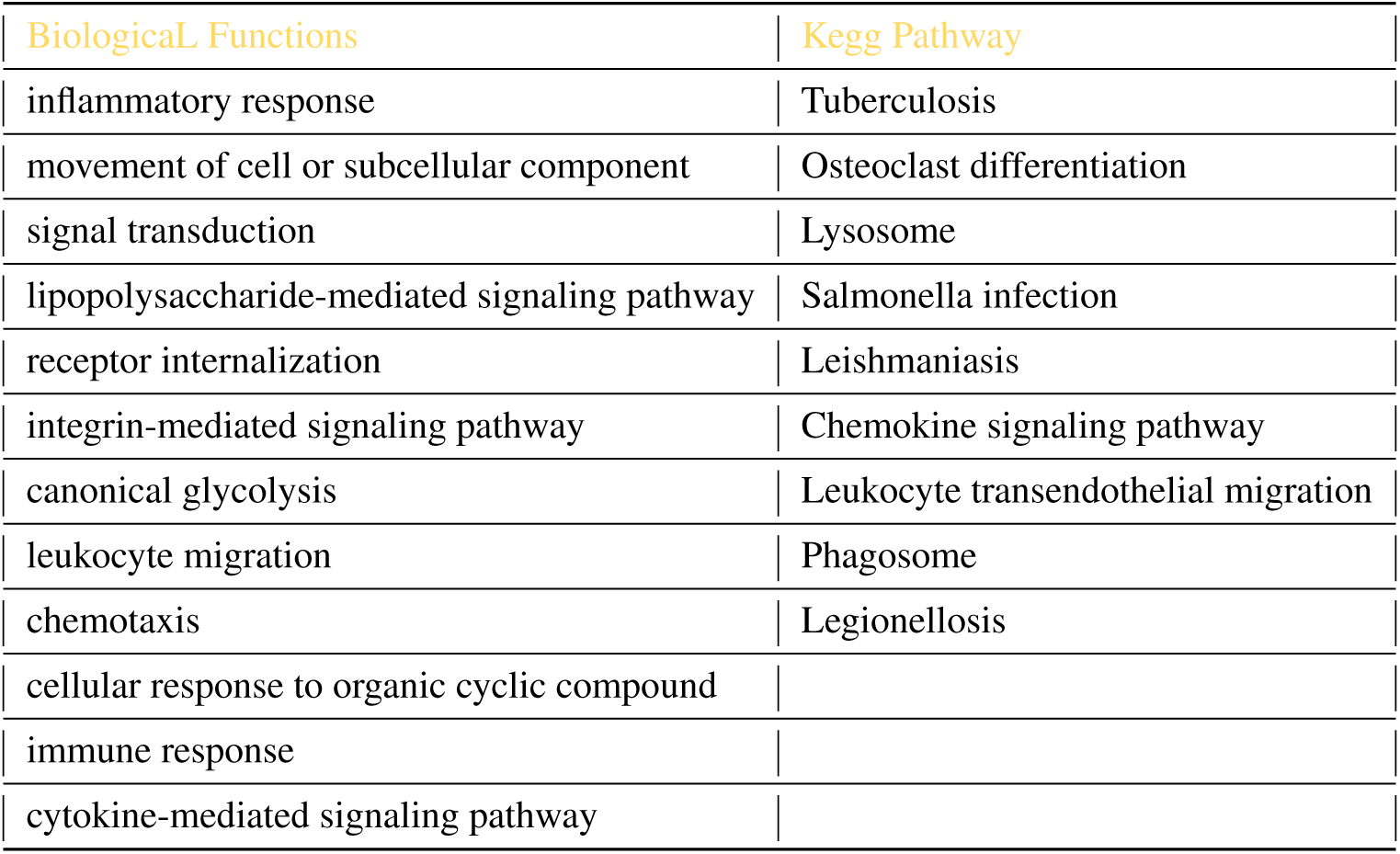
Biological function and KEGG pathway of module Turquoise

| **Biological Functions** | **Kegg Pathway** |
| --- | --- |
| inflammatory response | Tuberculosis |
| movement of cell or subcellular component | Osteoclast differentiation |
| signal transduction | Lysosome |
| lipopolysaccharide-mediated signaling pathway | Salmonella infection |
| receptor internalization | Leishmaniasis |
| integrin-mediated signaling pathway | Chemokine signaling pathway |
| canonical glycolysis | Leukocyte transendothelial migration |
| leukocyte migration | Phagosome |
| chemotaxis | Legionellosis |
| cellular response to organic cyclic compound |  |
| immune response |  |
| cytokine-mediated signaling pathway |  |

**Table 4.** Biological function and KEGG pathway of module blue

| Biological Process | KEGG Pathways |
| --- | --- |
| translational initiation | RNA transport |
| viral transcription | Ribosome |
| rRNA processing |  |
| nuclear-transcribed mRNA catabolic process, nonsense-mediated decay |  |
| transcription, DNA-templated |  |

#### 0.2.3 Co-expression Network

The Network of 2 modules one which contain RHBDD2 (Turquoise) and other which is negative correlated with RHBDD2 (Blue) shown in Figure7. In module turquoise green color was used to show the expression level in the disease. The dark green color is showing that the expression of a gene is high in disease while light green showing that the expression level of these genes is low in disease. The network of module blue which is negatively correlated with RHBDD2 and downregulated in disease is shown in the Figure. The color red is chosen here to show the expression level. Dark red nodes showing genes that have high expression in disease while light red shows the genes which have low expression in disease. The networks of Hub genes contain top 10 hub genes in the both network.

**Figure 7.**
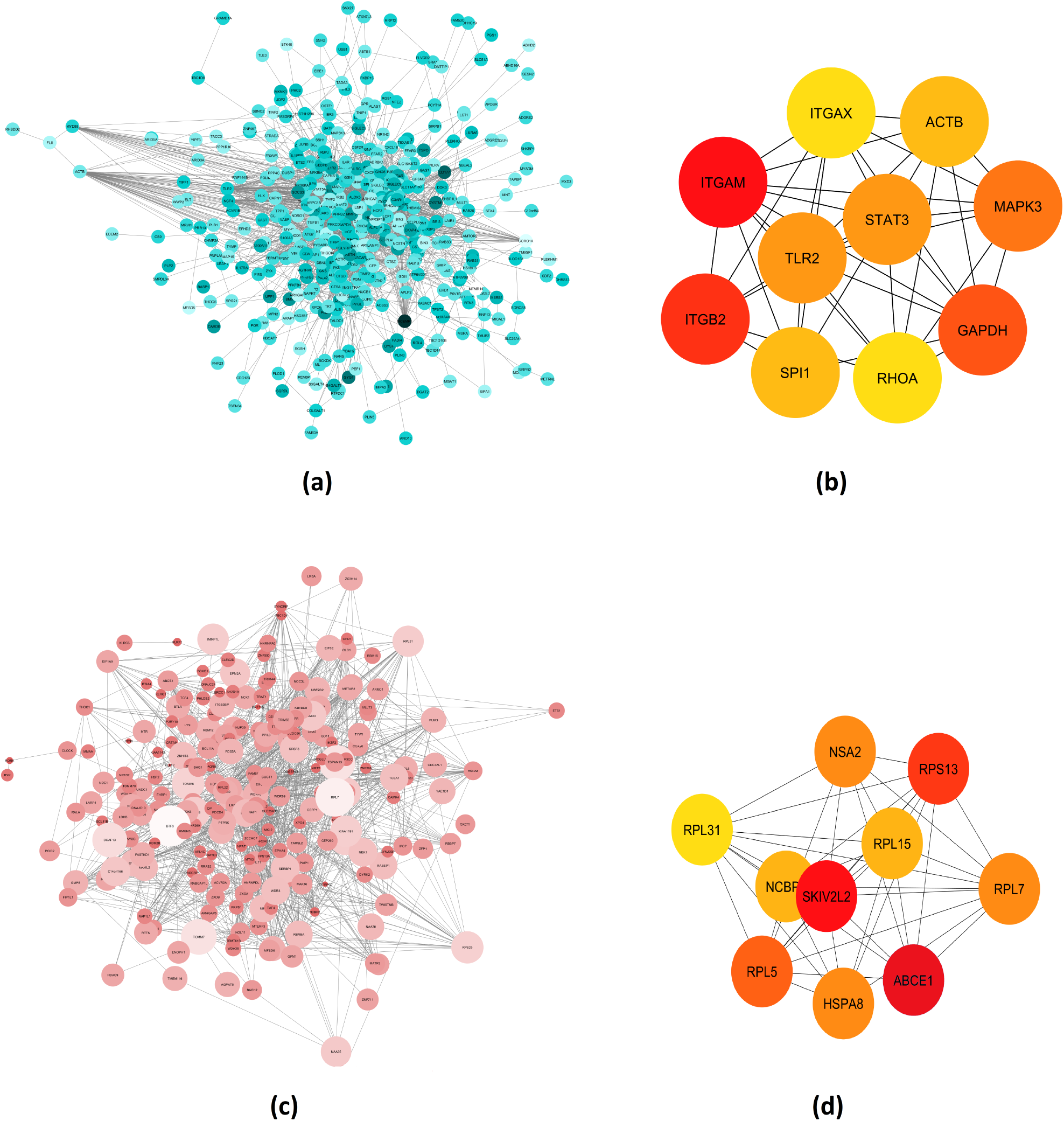
(a) representing the co-expression network of module Turquoise which contain RHBDD2 and the genes which are positively correlated with RHBDD2 and up-regulated in disease (b) Hub gens of module Turquoise(c) representing the co-expression network of module Blue which contain genes which are negatively correlated with RHBDD2 and down-regulated in disease (d) Hub genes of module Blue

**Figure 8.**
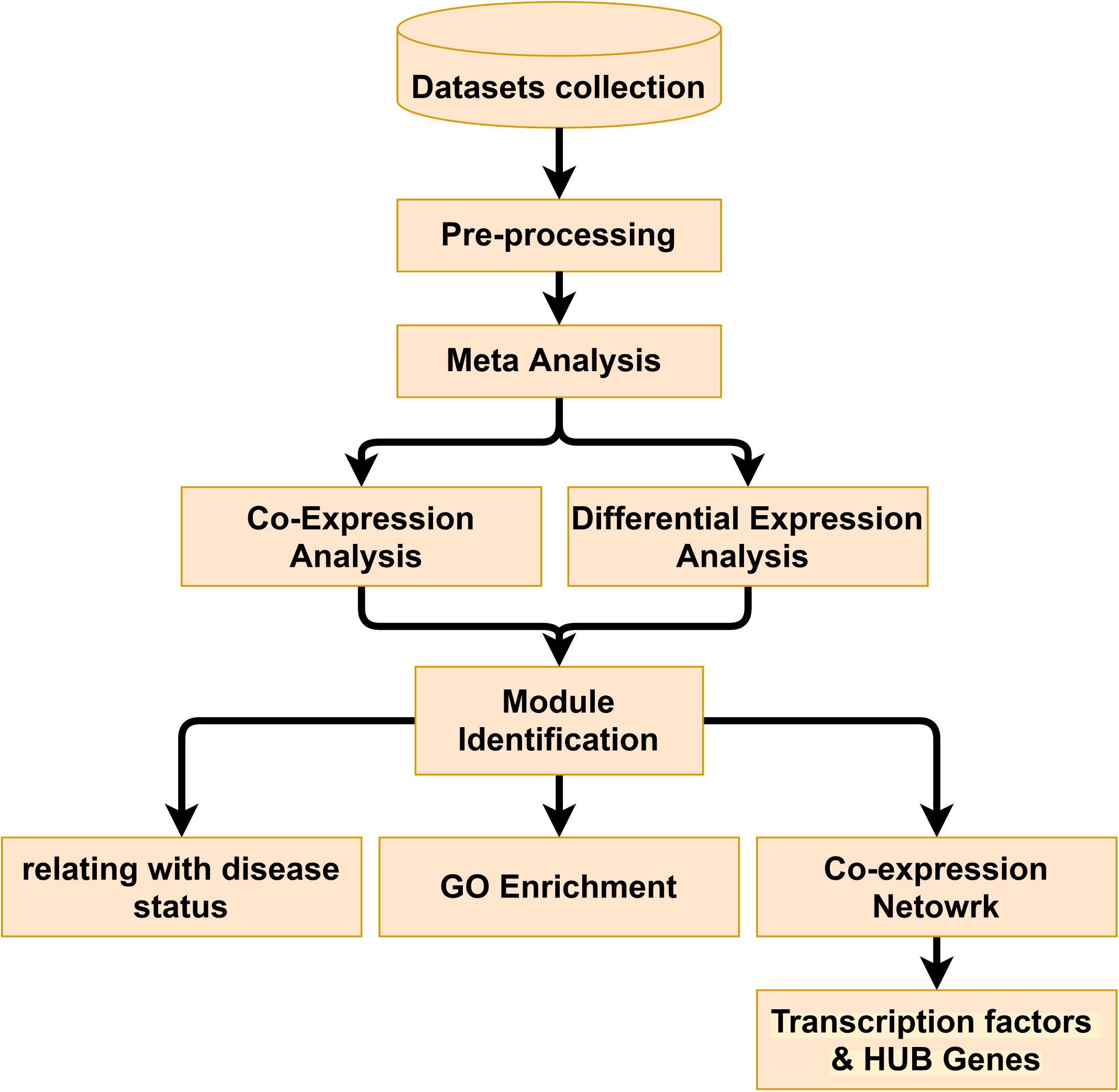
Flow of work (Methodology)

#### 0.2.4 Transcription factors

Transcription factors of module turquoise were SPI1, SATA5A, NFKB1, STAT3, MTA3, and JUND while transcription factors module Blue are MAX, MYC, and TAF1 all with AUC >0.5 and NES >3.

#### 0.2.5 Meta-Transcriptome Analysis

Meta transcriptome analysis revealed that the genes which are positively correlated with RHBDD2 are also up-regulated in disease and the genes which are negatively correlated with RHBDD2 are down-regulated in disease. Another Meta transcriptome analysis was done to check the relationship between RHBDD2 and Estrogen receptor which indicated that RHBBD2 has a negative relation with Estrogen receptor.

## Discussion

Sepsis is a serious health situation caused by uncontrolled infection and septic shock is a severe condition of sepsis. RHBDD2 is a member of the rhomboid superfamily which in a previous study found to overexpress in different types of cancer. Now, this study investigates that RHBDD2 is overexpressed in sepsis and septic shock too which is satisfy by limma and GeneMeta. The study reveals that most of the significant genes are co-expressed along with RHBDD2. These genes are regularly mentioned in literature, in relation to Sepsis and Septic shock.In addition, few of these genes are further mentioned as biomarkers. 3 co-expression modules obtained after WGCNA aalysis of RHBDD2-correlated genes, among which two modules (module “blue”, module “turquoise) were significantly associated with disease and were potential sepsis-associated modules. The enrichment analysis of module turquoise indicated that biological functions, such as; “receptor internalization”, “integrin-mediated signaling pathway”, ‘canonical glycolysis’, ‘leukocyte migration’, ‘chemotaxis’ and ‘inflammatory response’ are important factors in sepsis deregulation^16,17^. The immune response in sepsis the immune system plays a fundamental role in sepsis and septic shock. Cytokine increases the ER stress in sepsis and septic shock RHBDD2 which have an association with ER stress marker in cancer^18,19^. The module blue which carries genes that are down-regulated in sepsis negatively correlated genes with RHBDD2 involved in “translational initiation”, “viral transcription”, “nuclear-transcribed mRNA catabolic process” and “rRNA processing”. Sepsis produces a comparatively specific resistance in skeletal muscles that damages the capacity of this amino acid to stimulate protein synthesis and translation initiation^20^. Viral transcription is the consequence of viral sepsis^21^. RHBDD2 containing module showed the involvement of ‘Tuberculosis pathway’, which is an infrequent complication of tuberculosis, and is related to septic shock with multiple organ dysfunction^22–24^. ‘Osteoclast differentiation and ‘Lysosome pathways’ ware upregulated, which are implicated in septic shock and disregulation in latter causes exacerbate human sepsis^25,26^. Pathways of “RNA transport” and “Ribosome” of module blue were also found to associate with sepsis. All the pathways and biological functions of module turquoise found to be connected with sepsis and septic shock while module blue is found to be involved in basic cellular function and effect in the ribosomal pathway.

Through network analysis, 10 hub genes were selected using Cytohubba from two RHBDD2-sepsis-associated co-expression modules, which included hub genes, known to play role in sepsis and septic shock and their related pathways. The hub genes TLR2 and TLR4 are key genes in sepsis and septic shock and associate with module contain RHBDD2. TLR2, linked with cardiac dysfunction, deleterious systemic inflammation, and acute kidney injury, acts synergistically in sepsis and septic shock^27,28^. Mitogen-Activated Protein Kinase-3 (MAPK3) is a high differential expressed gene and a hub gene in sepsis and septic shock^29^. ACTB and GAPDH work as housekeeping genes^30^. Other hub genes RHOA, ITGAM, ITGB2, RvD2 and ITGAX also play an imperative role in sepsis and septic shock^29,31–34^. RHBDD2 was not found as a hub gene but it has interaction with FLII and then FLII interact with ACTB the hub gene. FLII appears in turquoise module, indicating a potential link between RHBDD2 and ACTB. RHBDD2 and its co-expressed hub genes and their involvement provides a piece of initial evidence that RHBDD2 is also an important factor in sepsis and septic shock, as it is regulated with its co-expressed genes and their respective roles are discussed above. The hub genes of module blue also reported in sepsis and septic shock. ABCE1, RPL15, RPL5, RPS13, RPL31, and RPL7 involved in protein binding and ribosomal functions^35,36^. NCBP2 levels reduced over the passage of sepsis. HSPA8 and SKIV2L2 are also functional importance in sepsis^37,38^.

Nine Transcription factors were the result of iRegulon. TLR4 signaling pathways that lead to NF-kB activation which plays a fundamental role in the pathophysiology of sepsis and septic shock and inhibition of NF-kB activation is capable to correct the cardiovascular functional abnormalities^39^. Transcription factors (SPI1 and the SATA5A) play roles in the regulation of RHBDD2 which also involved in sepsis^38,40,41^. So far there is increasing evidence that the RHBDD2 plays a part in sepsis. The Figure 5 (a) showing that RHBDD2 has an inverse relation with the Estrogen receptor in sepsis and septic shock as in cancer. Although the exact contributions of RHBDD2 to sepsis are not clear yet this study suggests that RHBDD2 a marker of sepsis and septic shock, further research is necessary as could be a potential transcriptomic marker for sepsis.

## 1 Methods

### 1.1 Dataset collection

In translational bioinformatics, microarray has its importance. Microarray provides high throughput genomic data which is used to evaluate thousands of genes at a one-time point^42,43^. It is used to identify the expression of genes that are expressed in different biological conditions^44^. To investigate the involvement in biological experimental conditions these genes are mapped to known annotation and regulation pathways based information. For the collection of microarray datasets, the publically available with MIAME (Minimum Information about a microarray experiment) rules and regulations^45^. Online repositories Gene expression Omnibus (GEO) and Array express where gene expression datasets of different biological conditions are available. We collect microarray gene expression Datasets for sepsis and septic shock from GEO and Array express Gene expression omnibus and Arrayexpress databases were thoroughly searched using the keywords ‘sepsis’, ‘septic shock’ AND human^46,47^. Datasets inclusion/exclusion criteria were defined as follows

1. Gene expression data based only on one of the most widely used Affymetrix platform AFFY HG U133 Plus 2 were selected to avoid intra platform biases.
2. Datasets having less than 12 samples were discarded to ensure enough representation from each population.
3. Only samples extracted from human blood/tissue were included in the cohort and cell line, mouse model-based data was excluded to avoid biological heterogeneity.
4. Only datasets having both normal, sepsis and septic shock samples were selected to assure representation of both groups from each population.
5. Datasets with available raw data i.e. cel files (having intensity of gene expression) were selected. Seven datasets with stages of sepsis) passed the filter and were used in the downstream analysis
6. Raw data comprising of .cel file for each sample in each dataset was downloaded. Bioconductor package Affy was used to read these cel files.

### 1.2 Pre-Processing

Preprocessing include three-step

1. Quality Check
2. Normalization
3. Reduction of probes

The quality of the datasets is most important to obtain more authenticated results. The biological and technical quality of each sample was evaluated using the following 5 parameters^48–53^. All these parameters were calculated using the yaqcaffy package by Bioconductor, and the samples appearing as outliers in more than three of these parameters (Glyceraldehyde 3-phosphate dehydrogenase (GAPDH),Beta Actin, Average Background Noise(AvBN), Percent Present(PP), Scaling factor(SF)) were removed from the Datasets and consequently from the downstream analysis.

#### 1.2.1 Normalisation

Normalisation was done using the MAS5 normalisation technique. Due to the simplicity of the algorithm behind MAS5 has performed well alongside multiarray methods^54^. “Affy” the Bioconductor package of R is used for MAS5 normalisation.Principal component analysis (PCA) plots were draw check the normalization

#### 1.2.2 Reduction of probes

A gene can have more than one probeset on the microarray chip, which may bring inconsistency or even contradictory results. To reduce the complexity of data, we used the Jetset algorithm to choose the best representative probe for each gene^55^. After preprocessing the next steps were Co-expression analysis and Differential expression

### 1.3 Meta-Analysis

#### 1.3.1 Differential expression enalysis

The differential expression of all genes was attained using Linear Models for Microarray Data (limma) [and meta differential expression was obtained by GeneMeta^56–59^.

#### 1.3.2 Co-expression analysis

Co-expression analysis is used in many studies to identify the functions of genes by making the module of genes that behave in a similar manner. Co-expression analysis as consisting of the following 4 steps^60^.

1. Similarity matrix
2. Module identification
3. Network construction
4. Go enrichment

#### 1.3.3 The similarity matrix

The co-expression network is based on the gene to the gene similarity matrix, which describes the expression pattern of genes in a group of samples^60^. For all seven datasets found the correlation of RHBDD2 with each gene using Pearson’s correlation was obtained^61,62^. Then Meta correlation of these correlations was obtained using R library “metacor”. After this, set a threshold on counts obtained by metacorrelation to select the top 1000 correlated genes with RHBDD2. These 1000 genes plus RHBDD2 then used for finding a combine similarity matrix. An algorithm was set, which take these 1001 genes and get the MAS5 normalized expression of all seven datasets of these 1001 genes and find the Meta correlation of these genes and form the similarity matrix^63^.

#### 1.3.4 Module identification

The weighted gene co-expression network analysis (WGCNA) is a data mining method used in systems biology which is based on pairwise gene co-relation. Co-relation networks that allow the generation of testable hypotheses from high throughput data. Many types of different biological problems and questions can be solved using WGCNA^64^. WGCNA was done using the R package “WGCNA”. WGCNA was used for the identification of a group of genes that are co-expressed with RHBDD2. The similarity matrix obtained from the previous step is converted to a signed adjacency matrix which is further used for module identification. The obtained modules will further have provided for Network construction and Go Enrichment.

#### 1.3.5 Network construction

The module obtains from WGCNA the module which contains RHBDD2 and co-expressed genes of RHBDD2 were given to “Cytoscape”. Cytoscape is a bioinformatics tool for the visualization and analysis of the network. STRING (database) was a plugin with Cytoscape to find the protein-protein interaction in a network form. This network was further used to obtain Hub genes. To find the hub genes this network further transfers to “cytohubba” ^65^. Cytohubba returns the top 10 hub genes.

#### 1.3.6 GO Enrichment

To investigate the genes attributes from the controlled vocabulary Gene Ontology (GO) term enrichment analysis was used. GO enrichment was done using “DAVID” which is a tool for functional classification^66,67^. The list of genes of each module obtained from WGCNA was given to DAVID from which biological process and KEGG pathways list was taken by setting the threshold of FDR<0.1 or P-value<0.05

#### 1.3.7 Transcription Factor

To find the transcription factor we used iRegulon the Cytoscape plugin for obtaining transcriptional factors and motifs^68^. Which takes the input of genes name and finds several transcriptional factors or the base of NES score and AUC.

## Author contributions statement

A.A. conceived the computational analysis, A.A., M.R and H.S. Analysis the results, A.H.and A.F. computational work. All authors reviewed the manuscript.

